# Differentiation and quality control of smooth muscle cells from human pluripotent stem cells via the neural crest lineage

**DOI:** 10.1101/2023.05.31.543049

**Authors:** Peter J. Holt, Hongorzul Davaapil, Deeti K. Shetty, Aishwarya G. Jacob, Sanjay Sinha

## Abstract

The Sinha laboratory has developed protocols for differentiating human pluripotent stem cells (hPSCs) into vascular smooth muscle cells along developmental lineage-specific pathways. In development, paraxial mesoderm (PM), lateral plate mesoderm (LM) and neural crest (NC) linages each give rise to smooth muscle cells significant in a location-specific manner. Induced PSCs derived from patients enduring disease provide a platform from which disease-relevant cell models can be established in the laboratory. Here we describe a robust protocol for differentiating hPSCs into vascular smooth muscle cells via a neural crest lineage and the control steps required to ensure consistently high-quality differentiated cells.

## Introduction

Studying vascular smooth muscle cells (SMCs) *in vivo*, is usually limited to whole organ imaging such as echocardiography and magnetic resonance imaging. Investigating the cell biology of the vascular media is particularly challenging due to poor tissue access and availability. Post-operative procurement of primary tissue or cells to work with, especially healthy/control material, is equally difficult. Primary cell numbers are typically low, have low proliferative capacity and become rapidly senescent in culture, making this an undesirable approach for investigating molecular mechanisms. Non-human model organisms can provide *in vivo* insight but do not necessarily recapitulate human mechanisms. Consequently, there is a strong case for human *in vitro* methods for research purposes.

The ability to reprogram (’induce’) dermal fibroblasts or peripheral blood mononucleocytes into pluripotent stem cells (iPSCs) which can then be differentiated into the cell type of choice provides a suitable alternative to *in vivo* analysis. *In vitro* modelling using iPSCs provides opportunities to study many non-syndromic single gene mutations which impact many cell types and tissues (Brooks et al., 2022). Differentiation of human patient-derived iPSCs to obtain SMCs offers an ideal way to study e.g. aortic disease (Davaapil et al., 2020). IPSC differentiation is able to recapitulate key developmental stages as well as generating different cell types for study. This ‘disease-in-a-dish’ approach is especially useful for genetic disorders (Tiscornia et al., 2011), of which there are many that affect major blood vessels such as the aorta, even in young people (Chou et al., 2023; Cury et al., 2013). Diseases where single mutations are causal offer the opportunity to correct pathogenic mutations in patient-derived iPSCs (using for example CRISPR-Cas9) to provide isogenic control cell lines. Some diseases that lead to aortic problems include Marfan Syndrome, Ehlers-Danlos Syndrome and Loeys-Dietz syndrome.

SMCs maintain vascular tone to regulate blood flow and contribute to the extracellular matrix of the medial layer, which together provide much of the structure and elasticity or stiffness of the vessel. SMCs are known to play a major role in aortic diseases (Galitito J et al., 2017; Oller et al., 2017; Rombouts et al., 2022). All three major developmental lineages (paraxial mesoderm, lateral plate mesoderm and neural crest), give rise to smooth muscle cells that contribute to the medial layers of blood vessels. These contributions are location-specific, with different locations being specifically vulnerable in different diseases, e.g. the ascending aorta and aortic arch in Marfan Syndrome, regions which comprise smooth muscle cells of neural crest (NC) lineage (MacFarlane et al., 2019; Majesky, 2007). Consequently, studying disease-related SMCs *in vitro* requires differentiation methods that derive lineage-specific cells. A further caveat is that stem cell differentiations can be variable between laboratories or even for individual researchers (Volpato et al., 2018a) and that care must be taken in deriving SMCs in vitro (Sivaraman et al., 2021).

Here we describe differentiation of pluripotent stem cells into neural crest SMCs, building on previous work from our laboratory (Cheung et al., 2012, 2014; Granata et al., 2017; Serrano et al., 2019). We identify the important control points in the differentiation at which cells need to be characterised before continuing and the analytical steps required to do this. Without such characterisation, neither the lineage nor the final nature of the cells can be certain. Using quality control procedures as we describe ensures that the process is reproducible and generates high quality, lineage-specific SMCs consistently. Without this consistency, results of experiments using the cells will be variable and data interpretation may be confounded (Volpato et al., 2018b).

Identification of neural crest cells was made by detecting expression of p75 (p75 neurotrophin receptor, p75NTR; Nerve Growth Factor Receptor; NGFR) and Transcription Factor AP2 alpha (TFAP2A), which are identified with early neural crest (De Crozé et al., 2011a; Schorle et al., 1996; Zhang J et al., 1996), with TFAP2A indicating pre-migratory cells (George et al 2021) and early migratory cells (Calmont et al., 2009). Identification of SMCs was made by detecting expression of Transgelin (TAGLN; SM22a) and Calponin (CNN). The canonical marker for contractile vascular SMCs is the myosin heavy chain isoform, Myh11 (Miano et al., 1994). However, this marker is not readily observed in stem cell-derived SMCs *in vitro* (J. Tang et al., 2020), although low amounts can sometimes be detectable by immunocytochemistry (Guo et al., 2013). Alternatively, TAGLN and CNN have long been known as characteristic of smooth muscle cells (Samaha et al., 1996) and have an established use as markers of smooth muscle cells derived from stem cells in vitro (Alexander & Owens, 2012; Wang et al., 2011). TAGLN may even be involved in the vascularisation process (Tsuji-Tamura et al., 2021). Each of these proteins has three isoforms, all expressed highly in SMCs (Liu & Jin, 2016; Matsui et al., 2018). Commercially available antibodies are typically not isoform specific or have lower affinity when supposedly isoform specific, so we have used robust non-isoform-specific primary antibodies.

To determine the efficiency of differentiation we have used flow cytometry and immunostaining to detect NC and SMC makers, as these are standard techniques that are technically straight forward and relatively fast. qPCR could also be considered to meet such criteria, but detecting proteins rather than RNA expression is a more robust indicator of cell state.

## Materials & Methods

Full details of all reagents and media are provided in the Supplemental information.

### Stem cell culture & Maintenance

Pluripotent stem cells are stored in DMSO-free Stem Cell Banker under liquid nitrogen. All stem cell suspensions were manipulated very gently, to avoid disrupting colonies. ROCK inhibitor Y-27632 (ROCK-Y; 1 µl/ml; 10 µM) was used in E8c media when reseeding after passage until cells clearly attached well. This was achieved by the neural crest stage at the latest and was often attained whist maintaining stem cells on Vitronectin. Five commercially available wild-type (normal) human stem cell lines were used in this work: H9 (embryonic stem cell), HPSI0114-eipl_1, HPSI0214i-wibj_2, HPSI0314i-sojd_3 and HPSI0414i-seru_7 (all 4 hiPSC lines from the Sanger HipSci collection).

### Culture plate preparation

Vitronectin (VTN) coated culture plates were prepared by adding 240 µl Vitronectin-XF to 5.75 ml PBS. 1 ml of VTN/PBS was added per well of a 6 well plate (6WP) and incubated for at least 1 hour at room temperature before use. For storage over 24 h or more, 1 ml of PBS per well of a 6WP was added and plates stored at 37°C. Gelatin-coated culture plates (gel-MEF) were prepared by adding 0.1% gelatin in PBS to cover the base of the culture vessel and incubated for 1 h at room temperature. Liquid was removed and MEF medium added (1 ml per well of a 6WP) and placed in an incubator at 37°C overnight or until required. MEF feeder plates were prepared by seeding irradiated MEF cells onto gel-MEF plates. A vial irradiated MEF cells was thawed quickly and added to 13 ml warm MEF medium in a 15 ml tube and collected by centrifugation (270*g*; 3 min). Supernatant was removed carefully, pelleted cells resuspended in 9 ml MEF medium and transferred to feeder cells at 0.5 ml per well of a 6WP (18 wells/3x 6WP). Cells were left to attach overnight, medium changed and the cells used within seven days.

### Stem cells on feeders

Stem cells were thawed quickly and added to 13 ml warm KSRc in a 15 ml tube and collected by centrifugation (17*g*; 3 min). At this low *g*, the larger colonies pellet loosely at the bottom whilst smaller colonies remain in the supernatant. Supernatant was removed carefully, pelleted cells resuspended gently in KSRc with ROCK-Y and transferred to MEF cells. Typically, each vial of stem cells was resuspended in 1 ml medium and transferred to a single well of feeders in a 6WP with an additional 0.5 ml medium per well. Medium was changed daily. When colonies were sufficiently large and numerous, they were picked to VTN plates. Fresh media was added to harvested feeder wells to maintain the remaining colonies.

### Stem cells on Vitronectin

Stem cells banked after growth on VTN for several passages could be seeded directly to VTN plates rather than feeders. Stem cells were thawed quickly and added to 13 ml warm E8c in a 15 ml tube and collected by centrifugation (17*g*; 3 min). At this low *g*, the larger colonies pellet loosely at the bottom whilst smaller colonies remain in the supernatant. At this point, supernatant can be transferred to a new tube, cells collected by centrifugation (270*g*; 3 min), resuspended in KSRc with ROCK-Y and added to wells of a MEF feeder plate as a precautionary reserve. Once supernatant had been removed, pelleted cells were resuspended gently in E8c with ROCK-Y, and transferred to a VTN plate. Typically, 1 ml medium was used per initial vial of stem cells and transferred to a single well of a VTN 6WP with an additional 0.5 ml medium per well. Medium was changed daily, small colonies form from single cells at 3-4 days and can be passaged once colonies are 500 µm or so in diameter. When transferring stem cell colonies from MEF cells onto VTN plates, or between VTN plates when seeking to improve colony size and morphology, picking colonies is essential. Using a releasing agent and/or scraping cells to collect them will not be as effective in these circumstances. Once passaging on VTN was effective, cells on feeders were no longer maintained.

### Passaging stem cells on VTN

Medium was aspirated and each well washed with 2 ml PBS. 1ml ReleSR was added per well of a 6WP and left for 2 min. ReLeSR was aspirated and cells left for 4 min at room temperature (20-22°C; timing will vary for different temperatures). 1 ml E8c was added per well of a 6WP and left for 5 min. Plates were tapped on the side to encourage colonies to detach. When required, plates were left for a further 1-5 min. Once detached, colonies were collected to a 15 ml tube using a p1000 pipette very carefully (wide bore tips can be a help). Cells were collected and dispensed slowly to avoid breaking colonies. Using a p1000 pipette, 1 ml of E8c was used to wash any further colonies from the well and transferred carefully to the relevant tube. Cells were collected by centrifugation (17*g*, 1 min), supernatant removed and cells resuspended gently in 1 ml E8c (with ROCK-Y if required). Using a p1000 pipette, cells were added a few drops at a time to fresh wells in a VTN plate containing 1 ml E8c (with ROCK Y if required). After adding 2-3 drops (corresponding to 50-100 μL, roughly 1/10^th^ of resuspended cells), wells were checked with a microscope to see if the required density was present. If not, more cell colonies were added until the required density was attained. Fresh medium was added to original, harvested wells to allow remaining colonies to recover and regrown. These were retained until the passaged cells were doing well. If colonies did not detach properly from the plate, the medium was removed, cells washed gently with 2 ml PBS, fresh medium added cells left overnight before trying again.

### Picking clones/colonies

Collagenase and/or dispase can be used to liberate stem cell colonies from feeders, but will result in collecting all iPSCs, not just good-looking colonies. Colony picking is preferable and essential when seeking to improve colony size and morphology. Colonies for picking were marked on the underside of the culture plate using a Nikon object marker on an Olympus CKX41 microscope and plates were transferred to an MSC. A p1000 pipette was used to pick each colony, first circling the colony and then aspirating whilst scraping across it. Colonies were added directly to the desired plate, into 1.5 ml medium with 1 µl/ml ROCK Y.

### Freezing iPSCS

Colonies were collected from VTN plates using ReLeSR as described. Colonies were resuspended carefully in 1 ml Stem Cell Banker, placed in a cryovial and then directly into a −80 freezer. After 2 days to a week, vials of frozen stem cells were transferred to liquid nitrogen for storage.

### Differentiation of stem cells into neural crest smooth muscle cells

#### Stem cells to neural crest

To obtain colonies of a suitable size for differentiation, stem cells were passaged 24-28 h before starting differentiation. Typically, a colony size of approximately 1-2 mm in diameter and a density of 2-4 colonies per cm^2^ was used. With larger colonies, cells in the centre are less accessible to the factors driving differentiation. If colonies are too small, they may not retain stem-ness well and may not reattach well when passaged.

#### Passaging cells using TrypLE

Neural crest and smooth muscle cells were passaged using TrypLE, since single cell suspensions were desired. Medium was removed from each well, cells washed with 2 ml PBS and 0.5 ml TrypLE added to each well. Cells were placed at 37°C and checked regularly until nearly all cells had detached and were in suspension (typically 3-5 min, but could be up to 10 min for confluent mature smooth muscle cells). Cells were transferred to a 15 ml tube, collected by centrifugation (17*g*, 3 min) and resuspended in fresh medium (typically 1 ml per well of a 6WP harvested). Cells were seeded at 0.3-0.4 x10^6^ cells per well in a fresh gel-MEF 6WP.

#### Neural crest to smooth muscle cells

24-48 h after passaging stem cells on VTN, E8c medium was exchanged for FSB medium (1.5 ml per well of a 6WP). FSB medium was refreshed each day for three further days (d0-3). On d3, cells were passaged as single cells using TrypLE onto gel-MEF plates; this is neural crest passage 1 (NC p1). Cells were fed daily with FSB and passaged when cells approached confluency, seeding at 0.3-0.4 x10^6^ cells per well in a new gel-MEF 6WP. From NCp4 onwards, excess cells were banked; neural crest cells tolerate banking well. This means that cells can be expanded and banked at early NC passages for future work on cells from the same differentiation. Also, if working with cell lines that have different proliferation rates, the faster-growing lines can be banked until all lines have reached the required stage.

#### Neural crest to smooth muscle cells (SMCs)

From NC p3 onwards, cells were differentiated into smooth muscle cells. For each neural crest stage differentiated into SMCs, some neural crest cells were also passaged to maintain neural crest. For differentiation, FSB medium was removed and PT medium added. Medium was renewed after 24 h and hence every 48 h for a total of 12 days (PT d12). Cells were passaged at 80-100% confluency and reseeded at maintain 50% confluency in the new wells, typically 0.4-0.5 x 10^6^ cells per well in a fresh gel-MEF 6WP. Smooth muscle cells thrive on cell-cell contacts and do not do well as single, sparsely distributed cells. Maintaining good confluency for the later days in PT is especially important and passaging at PT d8-d12 was avoided as far as possible.

#### Smooth muscle cell maturation

At PT d12, medium was changed to MEF medium, which was renewed every 8 h. SMCs were typically used for analysis after maturation for 14 days in MEF medium.

### Analytical methods

#### Immunocytochemistry

Sufficient room temperature 4% PFA in PBS to cover the cells (e.g. 0.25 ml per well of a 12-wellplate) was added to wells of a tissue culture plate and incubated for 10 min. Fixed cells were washed twice with PBS, PBS added to the wells and plates stored at 4°C until needed.

Neural crest cells were permeabilised, and all antibody dilutions and incubations done, in buffer (PBS with 0.2% Saponin and 5% FBS). After permeabilisation cells were blocked with 20% FBS in PBS for 30min. Blocking buffer was and primary antibodies added (anti-TFAP2A, anti-p75). Primary antibody incubation was at either room temperature for 2 h or over-night at 4°C with gentle rocking.

Smooth muscle cells were permeabilised with 0.1% Triton X-100 in PBS for 10 min and then blocked with 5% BSA in PBS for 30 min. Antibody dilutions and incubations done in PBS with 1% BSA. Blocking buffer was removed and primary antibodies added (anti-CNN, anti-TAGLN). Primary antibody incubation was at either room temperature for 2 h or overnight at 4°C with gentle rocking.

Primary antibodies were removed and cells washed gently three times with PBS. Secondary antibodies were added and cells incubated at room temp for 1 h shielded from light. An equal volume of DAPI at 1/5000 was added and cells incubated for a further 10 min. Secondary antibodies and DAPI were removed and cells washed gently three times with PBS. PBS was added for short-term storage (up to a week), DABCO added for longer-term storage and samples kept at 4°C until required for imaging.

All images of immuno-stained cells were taken using a Leica DMI4000 microscope in the Jeffrey Cheah Biomedical Centre imaging core facility. Basic image processing was done using Fiji (ImageJ) and analysed using QuPath (analysis protocol available upon request).

#### Flow Cytometry

(Reagent and antibody details are given in the Supplementary Information & SI Table 8).

All flow cytometry was done using an Attune NxT flow cytometer in the Biomedical Research Centre-Phenotyping Hub at the Jeffrey Cheah Biomedical Centre and data processed using FlowJo.

Cells were collected using TrypLE and diluted with an equal volume of PBS. Cells were collected by centrifugation (300*g*, 3 min), washed with 5ml PBS and collected by centrifugation. Cells were resuspended in 250 µl room temperature (RT) 4% PFA/PBS and incubated for 10 min in a 1.5 ml tube. Cells were collected by centrifugation, the PFA removed and cells washed twice with PBS. Cells were resuspended in 1 ml PBE buffer (PBS with 0.5% BSA and 2 mM EDTA) and stored at 4°C until required.

Neural crest cells and SMCs were immune-labelled the same way, differing only in the antibodies used. Samples were mixed by vortex briefly. Cell samples were divided equally into 1.5ml tubes for unstained controls, isotype controls and test samples. Use low retention tubes to avoid fixed cells getting stuck to the sides while spinning. Cells for each were pelleted by centrifugation (2000*g*, 3 min) and washed with 200 µl of flow buffer (PBS with 0.2% saponin, 5% FBS and 2 mM EDTA). Cells were pelleted by centrifugation, supernatant removed, then isotype controls and test samples were resuspended in 40 µl buffer. Unstained controls were resuspended in 50 µl buffer. 5 µl of each of control antibodies, AF488 labelled and AF647 labelled, were added to the isotype controls. 5 µl of each of anti-p75/NGFR-AF647 and anti-TFAP2a-AF488 were added to neural crest cell test samples. 5 µl of each of anti-TAGLN-AF488 and anti-CNN-AF647 were added to smooth muscle cell test samples. All samples were mixed briefly by Vortex and incubated at room temperature for 2 h. All samples were pelleted by centrifugation, supernatant removed and then washed with 1 ml buffer followed by 1 ml PBE and finally resuspended in 1 ml PBE. Samples were kept at 4°C until analysed.

## Results & discussion

H9 embryonic stem cells were maintained in 6-well vitronectin-coated plates and from each of three different passages half the cells were differentiated to neural crest (NC), providing three differentiation replicates. For each of these replicates from NCp4 onwards, samples of cells were taken for analysis by flow cytometry and immunocytochemistry (immunostaining; ICC) whenever possible. For each of these replicates from NCp4 onwards cells were differentiated into smooth muscle cells (SMCs). The overall process is summarised in Fig.1. Whenever possible, samples of SMCs at PTd4, PTd8, PTd12 and serum d14 were taken for analysis by flow cytometry and ICC. In all, 21 NC samples were analysed by flow cytometry and 26 by ICC, spanning NCp3-12. For SMCs, 65 samples were analysed by flow cytometry and 60 by ICC.

**Figure 1.**
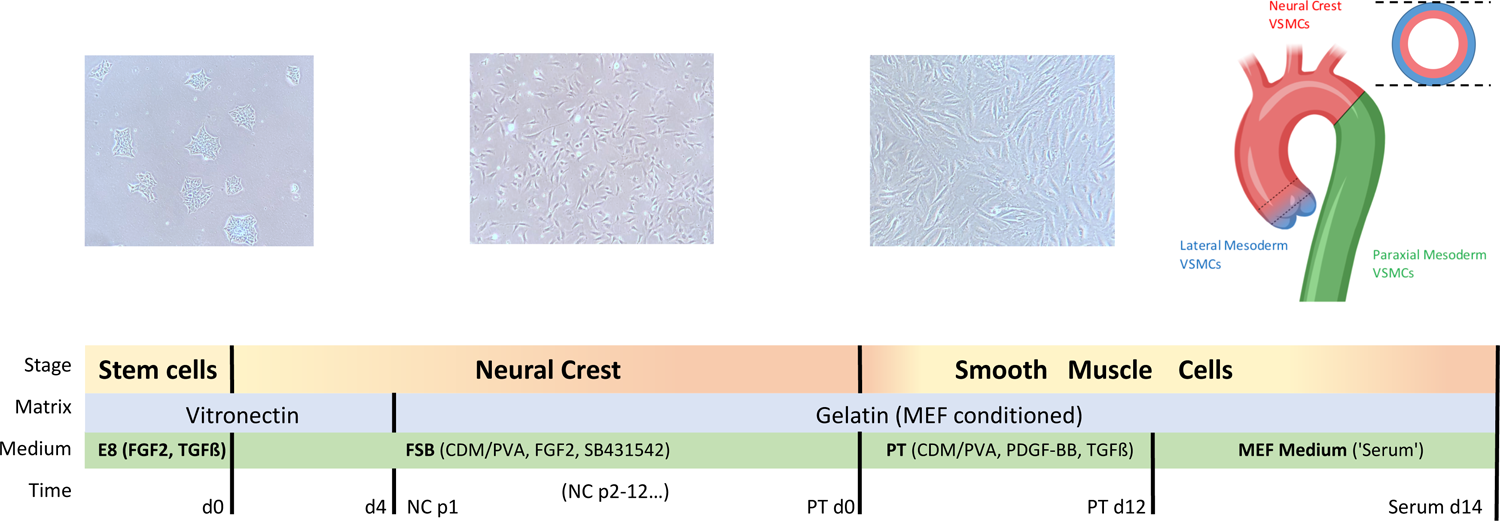
Differentiation schematic

### Stem cell culture and Neural Crest differentiation

Starting with a good stem cell culture is essential for any differentiation, and this work is no exception. Small colonies, which maintain stem cell colony character, whilst permitting thorough access for soluble factors, are the best starting point for differentiation. Typical stem cell cultures are illustrated in Fig.2A-D. Early stages show a mixture of colony sizes and single cells (Fig.2A). With careful passaging a more consistent colony size (about 400-800 µm) and few single cells can be achieved, suitable for differentiation into neural crest (Fig.2B). This is a first control point; if stem cell colonies are sub-optimal, then initial neural crest differentiation will be impaired.

**Figure 2.**
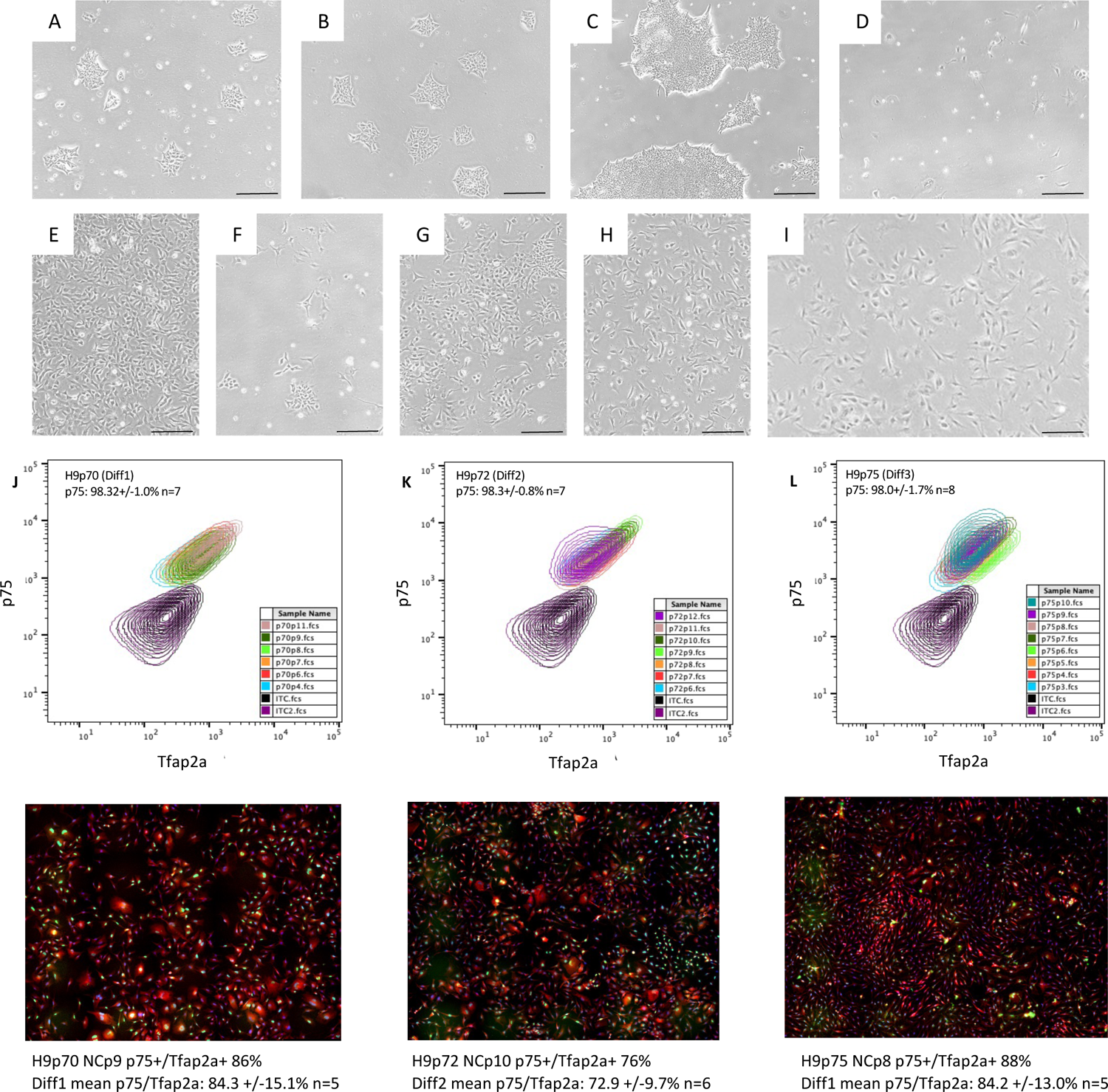
Differentiation of stem cells to Neural Crest. A: Typical stem cells. B: Stem cells good for differentiating to Neural Crest. C: Stem cell colonies too large and starting to merge, require passage before differentiating. D: Stem cells losing stemness and differentiating. E: Good NC p1-2. F: Poor NC p1-2. G: recovering NC p1-2. H: Good NC p3. I: Good NC density to add PT medium for differentiation into SMCs. Black scale bar indicates 200µm J: Differentiation 1 flow cytometry NCp4-10 and example ICC (NCp9). K: Differentiation 2 flow cytometry (NCp6-12) and example ICC (NCp9). L: Differentiation 3 flow cytometry (NCp3-12) and example ICC (NCp8). For immunostaining, TFAP2a id in green, p75 is in red, DAPI is in blue. White scale bar indicates 500µm.

We used ReLeSR for passaging stem cells as this gave small colonies consistently. Time in this reagent is critical, too short and cells do not detach well, too long and single cells become dominant. To passage, we incubated the cells for four minutes with the reagent, aspirated the solution and incubated for a further four minutes with the cells remaining coated with a thin layer of liquid). After this, an appropriate volume of medium was added to collect colonies. This strategy worked well here, but some optimisation may be needed for different cell lines. Stem cells should be handled gently during passaging and trituration kept to a minimum, although gentle trituration is sometimes useful for breaking larger colonies. Careful use of TrypLE or EDTA will work also, but require more care to obtain consistently small colonies suitable for differentiation. TrypLE treatment especially is prone to generate more single cells. EDTA often requires scraping to help detach colonies and colony sizes are more variable. Cultures with colonies that grow large and start coalescing are not suitable for differentiation (Fig.2C), as cells towards the centre of large colonies do not receive effective exposure to the factors required to drive the change. Stem cell cultures where colonies are too small and/or too many single cells occur can quickly show signs of differentiation (Fig.2D), changing morphology and exhibiting significant cell processes. In such cases, any suitable colonies should be picked to new culture wells and expanded until ready for differentiation.

FSB medium was added to stem cells (d0) and media changed daily for four days before passaging (d4) to give NC p1 cells. NC cells have a characteristic appearance, appearing as small, roughly triangular cells with short processes from the ‘corners’ and a pale halo on brightfield microscopy (Fig.2E-I). Four days before first passage is the guideline; if cells become confluent, passaging is better than letting them overgrow. If the initial passage does not go well and few cells attach, these cells typically proliferate slowly (Fig.2F). Feeding daily and being patient usually yields cells that proliferate and improve sufficiently for passaging (Fig.2G) and then recover from NCp3 onwards (Fig.2H). Maintaining cells at about 40% confluency when passaging gives good cell morphology and proliferation. When seeded too sparsely, these attributes are impaired. Differentiation into SMCs should be started at least 24h post-passage, at a density of 40-50% confluence (Fig.2I). From NCp3 onwards, cells required passaging every 2-3 days. This is a second control point; neural crest cells at p3 should look correct by brightfield microscopy or further passages may not be suitable for differentiation into smooth muscle cells.

Cultures at NCp1-2 were not assessed for NC markers, as these stages retain undifferentiated cells and non-specific neuroectoderm. Also, at NCp1-3, cells had rarely proliferated sufficiently for samples to be taken for analysis. However, if sufficient cells are available at NC p3, these should be tested for NC markers. From NCp4 onwards (exceptionally NCp3), samples were taken for analysis by flow cytometry and immunocytochemistry (immunostaining; ICC). For ICC to be effective, both in terms of staining efficiency and cell counting capacity, cell density needs to approach, but not reach confluency. Evenly distributed cells at a density of c. 70% works well (SI Fig.1). The results are summarised in Figure 2J-L. Flow cytometry revealed 97-99% cells positive for the NC marker p75 and ICC showed high proportions of cells positive for both p75 and TFAP2A: 84.3+/-15.1%, 72.9+/- 9.7% and 84.2+/-13.0% for the three replicates. These two markers can be challenging to stain for simultaneously; p75 is a surface marker, thus sensitive to detergents that permeabilise the cell membrane, and TFAP2A is a transcription factor located in the nucleus. ICC proved more robust than flow cytometry in this regard. This is almost certainly a function of using directly conjugated primary antibodies for flow cytometry as compared to primary with secondary antibody combinations for ICC, where signal is amplified. It may be possible to optimise these procedures; for example, there are very many detergents that could be tried as permeabilisation agents, some of which could offer improved performance (less disruptive to surface p75 whilst permeabilising both the cell and nuclear membranes effectively) than the saponin used here. (There is a large body of work in the protein structure and function field illustrating how different proteins can be uniquely responsive to even superficially similar detergents in a poorly predictable manner). Especially if this proved the case, then a primary/secondary antibody staining process may provide better resolution on flow cytometry. Sequential staining for p75 and then TFAP2A, with primary antibody incubations overnight, could also improve staining. Such variations were discounted for this work, though, as the process would become too lengthy for the purpose of establishing whether or not rapidly-growing cells should be differentiated.

Cultures showed an average of 70% or more neural crest cells from NCp4 onwards (Fig.4A). It is possible that the time spent in FSB medium in reaching p4 is significant, as well as the number of passages. However, these two factors are difficult to separate as cells need to maintained at a density of 40%< confluence, thence requiring regular passaging. It is likely that both time in FSB medium and selection/purification by passage are required, but this is not proven. As such, we recommend differentiating into SMCs from NCp4, provided that cells are > 90% p75+ positive and > 70% p75+/TFAP2A+.

### Smooth muscle cell differentiation

Smooth muscle cell markers TAGLN/CNN were determined by ICC and flow cytometry regularly through the 12 days in PT and at serum d14 (Fig.3). Concordance between the data from the two techniques improves from PTd4 to PTd12 and serum d14, the co-stained cell mean percentages being: PTd4 flow 70.4+/-10.3% and ICC 90.0+/-3.1%; PTd8 flow 71.9+/-11.9% and ICC 86.4+/-6.5%; PTd12 flow 79.6+/-13.8% and ICC 82.3+/-6.8%; serum d14 flow 82.1+/-13.7% and ICC 86.0+/-7.2%. (Fig.4B). The markers used are both cytoskeletal, so a harsher detergent (Triton X-100) could be used to thoroughly permeabilise cells for both techniques. Smooth muscle cells do not thrive at low densities, so it is important to maintain a c.40%< confluence, especially during the 12 days in PT medium. During the first 4 days and final 2 days in PT cells passage readily, if density is maintained, but proliferation is often slow if passaged during days 5-10. Once in serum to mature, cells typically need passaging no more than once before day 14. if in doubt, don’t passage; smooth muscle cells thrive on density.

**Figure 3.**
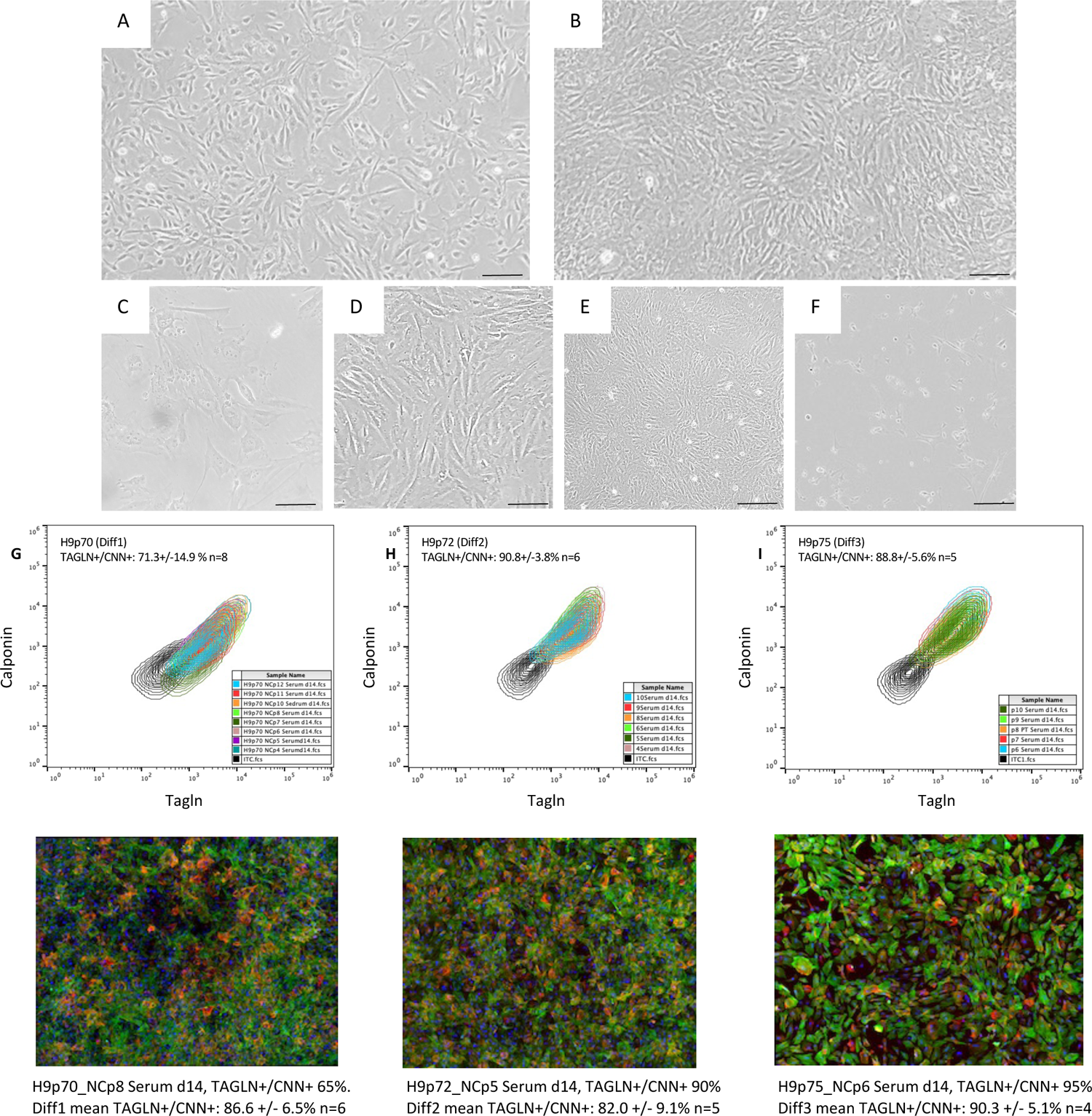
Differentiation of Neural Crest Cells to Smooth Muscle Cells. A: SMCs in PT at d12, 24h after passage. B: SMCs in PT at d12, confluent. C: Sparse SMCs in serum. D: Good density SMCs in serum. E: Over-dense SMCs in serum. F: SMCs in PT at d12 with some non-SMCs apparent. F: SMCs in serum with poor morphology and non-SMCs apparent. Black scale bar indicates 200µm. G: Differentiation 1 flow cytometry and example ICC. H: Differentiation 2 flow cytometry and example ICC. H: Differentiation 3 flow cytometry and example ICC. For immunostaining, TAGLN is in green, CNN1 is in red, DAPI is in blue. White scale bar indicates 500m.

The markers used here (TAGLN, CNN) are established markers of contractile cells and work well in flow cytometry and ICC quality control assays. Other possibilities include F-actin, which stains readily with Phalloidin conjugates and Smoothelin (Van Der Loop et al., 1997; van Eys et al., 2007). However, if a truly contractile cell type is required, a contractility assay (Bogunovic et al., 2019; Steucke et al., 2015) should be used as the definitive assessment of final cell quality. In this regard, traction force microscopy may be considered the definitive technique currently (Ahmed et al., n.d.) Correlation between NC (all passages) and final SMC (serum d14) marker expression is shown in figure 4C. High levels of NC marker expression generally correlate with high levels of SMC markers, but both should be determined. If NC markers are low, then the lineage of the final SMCs will not be correct; if the SMC markers are low, then the cells cannot be considered SMCs. Using the criteria of progressing from neural crest only when these cells are p75 90%+ by flow cytometry and 70%< p75+/TFAP2A+ by ICC yields consistently good quality SMCs, as determined by TAGLN+/CNN+ expression (typically 80%+ by flow cytometry and ICC).

**Figure 4.**
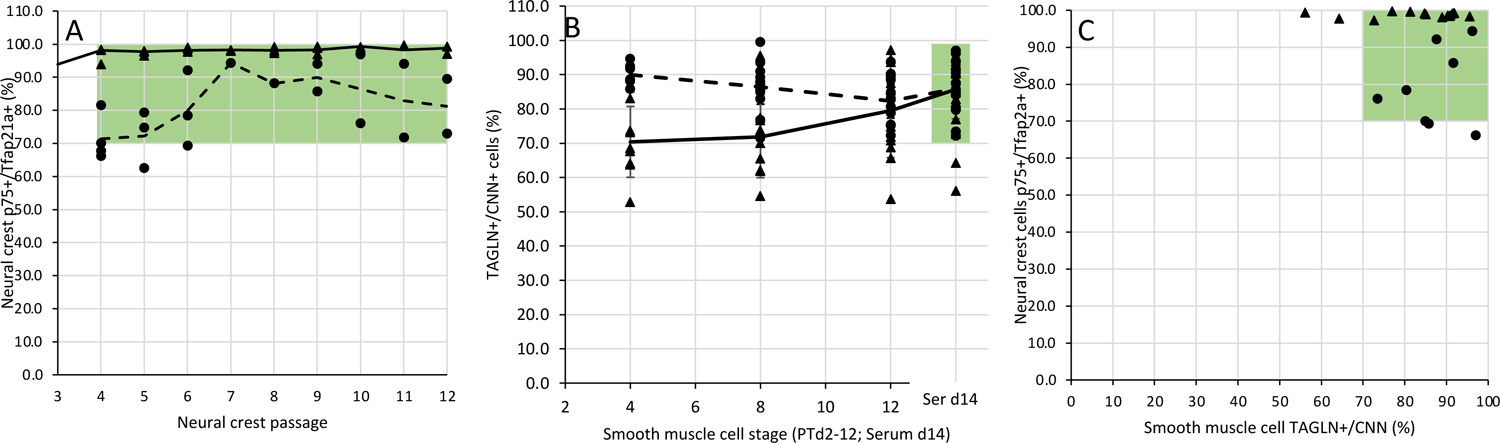
Differentiation progression. A: Neural crest marker expression over passages 4-12. Solid line indicates mean flow data, dashed line indicates mean ICC data. B: Smooth muscle cell marker expression over PTd4-12 and at serum d14. Solid black lines indicate mean flow data, dashed line indicates mean ICC data; error bars indicate +/-1 s.d. C. Correlation between neural crest marker expression (all passages) and final smooth muscle cell marker expression at serum d14. Flow data are plotted as triangles, ICC data as circles. Green shading indicates cell cultures that are 70%+ for the relevant markers at the various crucial neural crest stages (A) when differentiation to smooth muscle cells would start and for the final smooth muscle cells at serum d14 (B, C).

At all stages, proliferation can vary between lines, with some WT iPSC lines giving rise to robust cells that need passaging regularly, even in serum. Other cell lines, notably patient-derived disease lines, can be considerably more sensitive. This can only be determined empirically.

### iPSC differentiation

Having established the method with attendant quality control points using the embryonic H9 cell line, the NC SMC differentiation protocol was used on four wild-type iPSC lines. Samples from each were evaluated by ICC (Fig.5). For the four lines, TAGLN+/CNN+ cells levels were 83-92% for NC SMCs. The NC SMC protocol described here, with screening at the NC stages for cells expressing p75 and >70% expressing TFAP2A immediately before beginning differentiation into SMCs, thus proved robust and viable for both a standard embryonic stem cell line (H9) and four different iPSC lines. Precursors to this work have been successful in differentiating patient-derived iPSC lines for *in-vitro* disease modelling (Granata et al., 2017b; Serrano et al., 2019b). With the control measures we have now defined, consistently high-quality SMCs from a confirmed neural crest lineage should be obtained for such purposes. Provided that experiments begin with sufficient iPSCs, the ability to expand and bank cells at the neural crest stage means that multiple experiments can be done from cells of the same differentiation as and when required.

**Figure 5.**
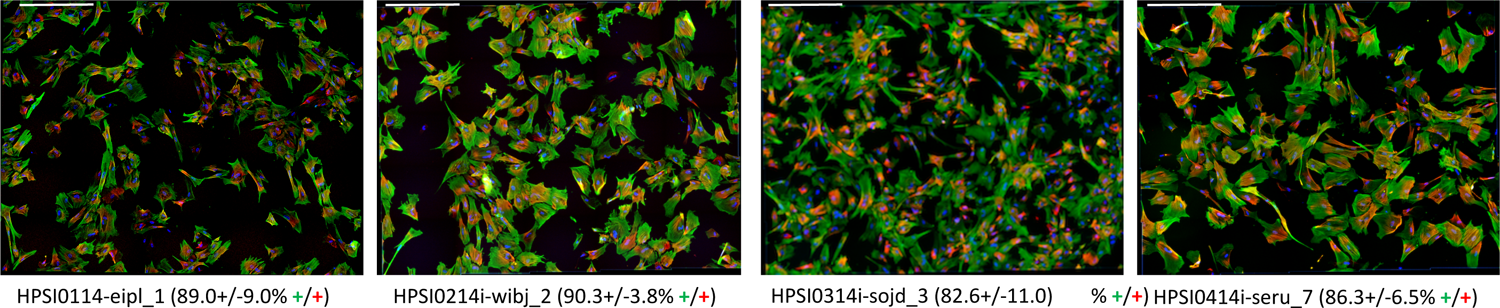
Differentiation of WT iPSC lines to Neural Crest Smooth Muscle Cells. Immunostaining of neural crest smooth muscle cells from differentiation of four WT iPSC lines. TAGLN is in green, CNN is in red and DAPI is in blue. Scale bar indicates 500µm. n=3 in all cases.

## Summary

This work describes differentiation of PSCs to SMCs via a neural crest lineage, for both a widely-used commercial human embryonic stem cell line and multiple human iPSC lines. A limitation to this study was that although care was taken to have a good representation across the process of differentiation, we were unable to collect samples for all analyses at all points for all replicates during the differentiation. Preference was given to collecting at least one sample for every point possible rather than replicates of any given point. However, analysing cell samples at key points, namely neural crest immediately before differentiation to smooth muscle cells begins and at the final smooth muscle cell stage, is essential to obtaining consistent results (Fig.6). Protocols for doing this are described in detail and data show clearly that our differentiation method is robust and reproducible when cells from neural crest p4 onwards are used.

**Figure 6.**
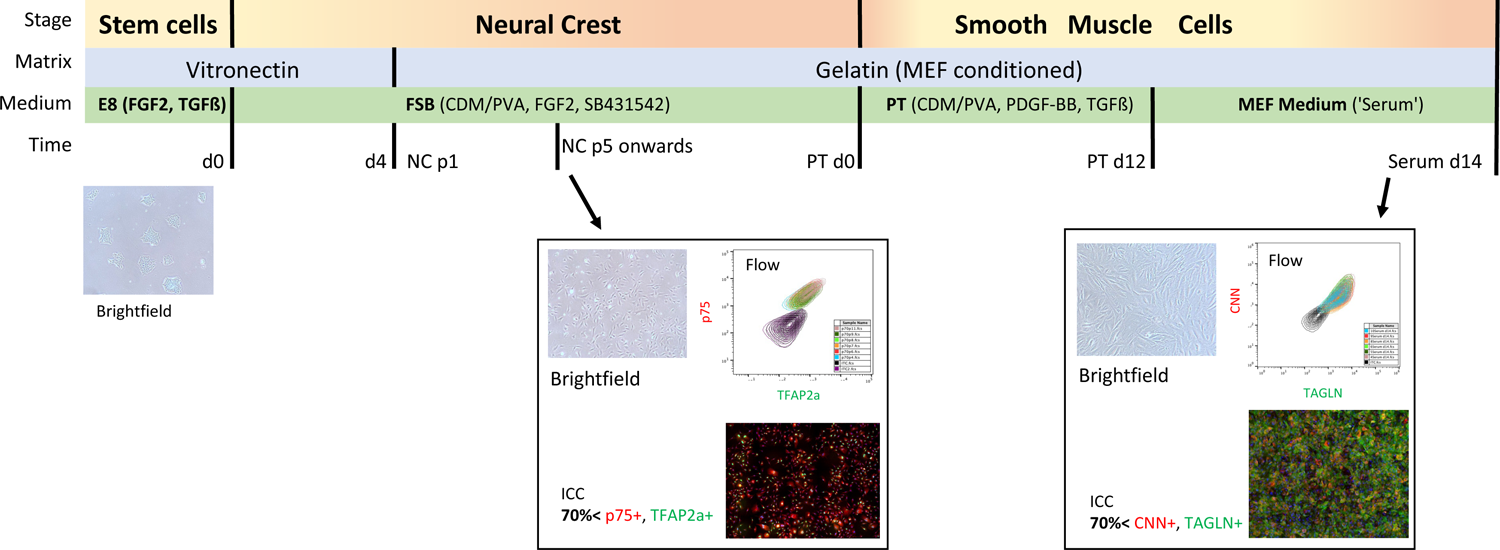
Differentiation schematic with quality control points.

It is notable that the neural crest in development comprises highly diverse cell populations that are both transitory and migratory (Bhattacharya et al., 2021; Dinsmore & Soriano, 2022; Kirby & Waldo, 1995; Schussler et al., 2021; Simões-Costa & Bronner, 2015). In mouse, early commitment to specific migratory routes and ultimate cell fate has been described (Chen et al., 2021). Expression of p75 and TFAP2A, which are identified with early migratory neural crest (De Crozé et al., 2011b), were used as markers in this work. The protocol described here generates neural crest cells exhibiting consistent levels of these two key markers over multiple passages (until at least p12). These cells are routinely stored under liquid nitrogen for future use without detriment (data not shown). However, identifying reliable characteristics of neural crest cells that migrate to specific cardiac destinations, notably to the great vessels, is desirable. Such knowledge could guide *in vitro* differentiation of a cardiac-specific neural crest lineage smooth muscle cells. It is likely that controlled exposure to various defined ratios of BMP4/Wnt1 is needed to drive such specific sub-lineages (Hong et al., 2022).

SMCs are a major component of the aortic wall and are largely responsible for generating the extra cellular matrix that provides much of the structure and elasticity or stiffness of the vessel. There is a significant genetic basis for aortic disease (Brownstein et al., 2017; Pinard et al., 2019) and smooth muscle cells (SMCs) are known to play a key role in aortopathies (Galitito J et al., 2017; Oller et al., 2017; Rombouts et al., 2022), although endothelial cells have been implicated too (Chung et al., 2007). Since the ECM has a significant role in vascular disease, thoroughly characterising ECM in health and disease could guide development of a protocol for maturing in vitro differentiated SMCs that deposit ECMs matching these states. Amongst vascular diseases where smooth muscle cells have a significant role, those affecting the aorta are particularly notable due to their severity and often acute manifestation. Aortic aneurysm is a significant and increasing cause of death world-wide (Bossone & Eagle, 2021; Lim et al., 2012). Treatment of susceptible individuals is limited currently to medication and lifestyle adaptations that limit blood pressure raises, with often urgent surgical intervention required to deal with aortic dilation, dissection or rupture (John Pepper, 2020). Areas that may prove interesting to study include the length of time between passages, the length of time SMCs spend in the final maturation medium and the effect these times have on ECM deposition and marker expression. iPSC-derived ‘disease-in-a-dish’ models are eminently suited to such work.

Neural crest lineage cells are also present in areas of the heart, and being able to identify and recapitulate such specific cell lineages *in vitro* could be significant in investigating mechanisms in non-vascular diseases (Erhardt et al., 2021).

Cardiac regeneration in organisms such as zebra fish (W. Tang et al., 2019) and mouse (Tamura et al., 2011) has been linked to cells of neural crest lineage, although a review of the work on mouse describes mixed results and it is merely theorised as possible in human (Erhardt & Wang, 2023). Consequently, the robust generation of cardiac neural crest cells *in vitro* may be useful in the cardiac regeneration field, but this currently remains highly speculative.

## Acknowledgements

This work was supported by the following British Heart Foundation grants: BHF Program grant (RG/17/5/32936, PJH, DS, HD and SS) and BHF Senior Fellowship (FS/18/46/33663, SS). This research was funded in whole or in part by Wellcome Trust (203151/Z/16/Z) and the UKRI Medical Research Council (MC_PC_17230). This research was supported by the Cambridge NIHR BRC Cell Phenotyping Hub. The authors gratefully acknowledge the Advanced Microscope Facility, JCBC, for their support and assistance in this work and in particular Darran Clements for his guidance.

## Author contributions

PJH conceived and performed experiments and analysis and wrote the paper. AJ, DS and HD differentiated iPSC lines. SS conceived and supervised the project. All authors reviewed the manuscript.

---

**Supplementary Figure 1.**
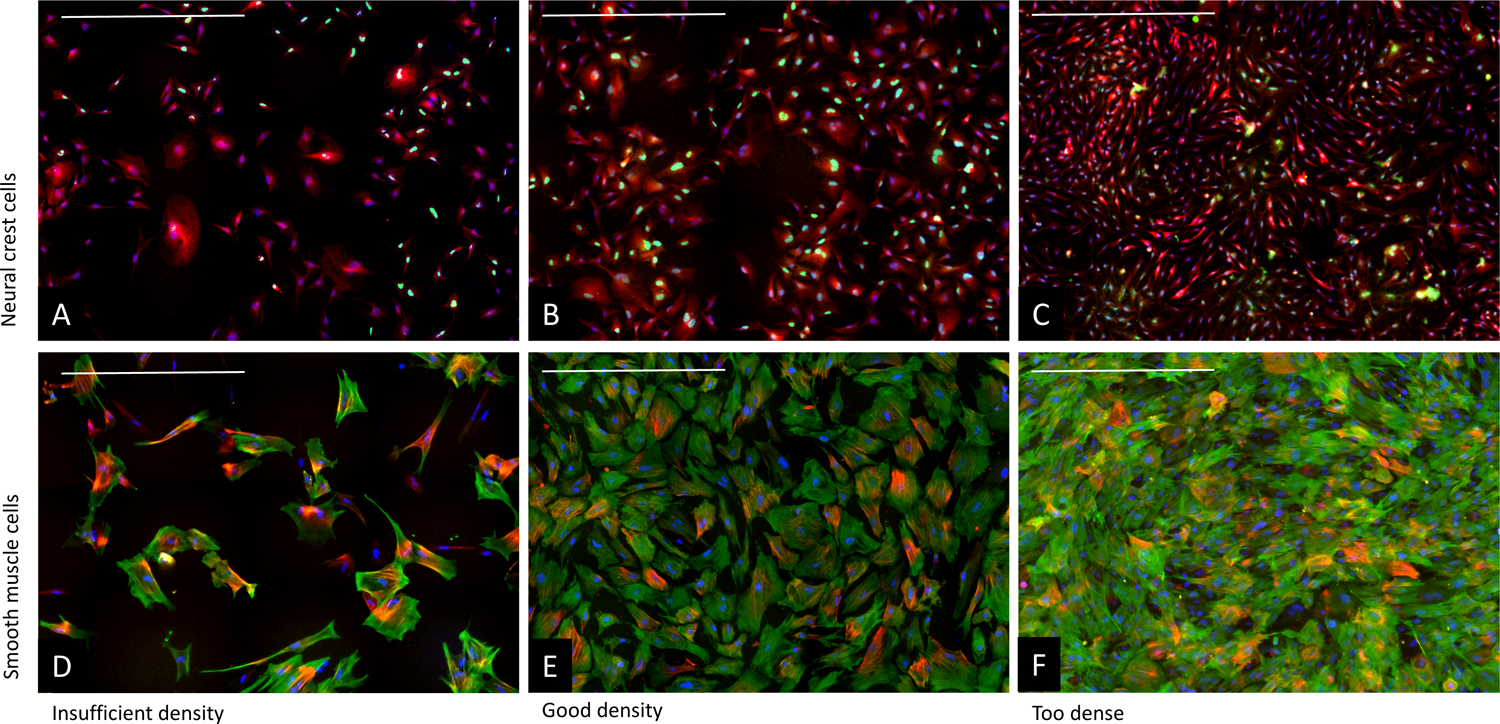
Cell density and ICC. Examples of Neural Crest and Smooth Muscle cells that are at insufficient density for effective counting (A, C), good density for staining and counting (B, D) and that are too dense for effective immunostaining and counting (C, E). Scale bar indicates 500µm.

**Supplemental Figure 2.**
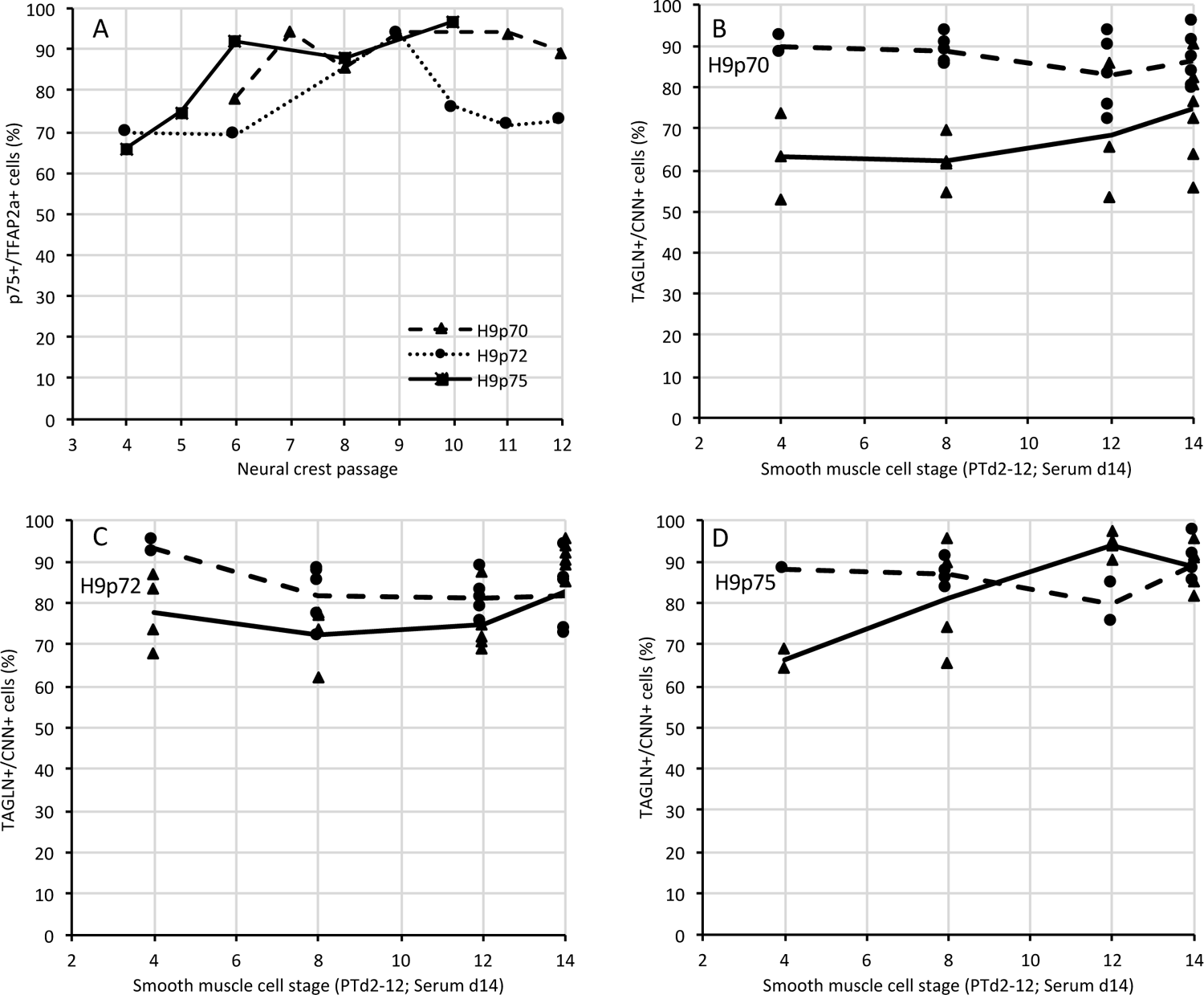
Differentiation progression. A: Neural crest marker expression over passages 4-12 for the individual differentiations, ICC data. B-D: Smooth muscle cell marker expression over PTd4-12 and at serum d14 for individual differentiations. Flow data are plotted as triangles (solid black line indicates the mean) and ICC data as circles (dashed line indicates the mean) for differentiations H9p70 (B), H9p72 (C) and H9p75 (D).

## Supplemental Information

**Supplemental table 1.** E8 medium.

| E8 medium |  |  |  |  |
| --- | --- | --- | --- | --- |
| Component | Source/product code | Stock concentration | Volume/amount | Final concentration |
| DMEM/F12 | Thermofisher 31330038 | N/A | 500 ml | N/A |
| Insulin-transferrin-selenium | Thermofisher 4140004 | N/A | 10 ml | N/A |
| Sodium bicarbonate | Merck/Sigma S8761-100ML | 7.5% | 5 ml | 0.075% (10 mM) |
| L-Ascorbic acid 2-phosphate | Merck/SigmaAldrich A8960-5G | 32 mg/ml (110 mM) | 1 ml | 60 µg/ml (0.22 mM) |
| Pen/Strep | Thermofisher 15140122 | 100x | 5 ml | 1x |
| Complete E8 medium (E8c): |  |  |  |  |
| TGF-β | Peprotech 100-21-100 | 2 µg/ml | 500 µl | 2 ng/ml |
| FGF-2 | Qkine Qk027 | 4 µg/ml | 3.125 ml | 12 ng/ml |

**Supplemental table 2.** CDM/PVA.

| CDM/PVA |  |  |  |  |
| --- | --- | --- | --- | --- |
| Component | Source/product code | Stock concentration | Volume/amount | Final concentration |
| Ham's F12 nutrient mix | Thermofisher 21765029 | N/A | 250 ml | N/A |
| IMDM | Thermofisher 21980032 | N/A | 250 ml | N/A |
| Concentrated lipids | Thermofisher 11905031 | 100x | 5 ml | 1x |
| Transferrin | R&D Systems 3188-AT-001G | 30 mg/ml | 250 µl | 15 mg/ml |
| PVA | Merck/SigmaAldrich P8136-250G | 10% | 5 ml | 0.1% |
| Insulin | Merck/SigmaAldrich 11376497001 | 10 mg/ml | 350 µl | 7 µg/ml |
| Pen/Strep | Thermofisher 15140122 | 100x | 5 ml | 1x |
| 1-thioglycerol (MTG) | Merck/SigmaAldrich M1645-25ML | (Neat) | 20 µl | (~0.5 mM) |

**Supplemental table 3.** PT medium.

| PT medium |  |  |  |  |
| --- | --- | --- | --- | --- |
| Component | Source/product code | Stock concentration | Volume/amount | Final concentration |
| CDM/PVA | N/A | N/A | 500 ml | N/A |
| PDGF-BB | Peprotech 100-14B-250 | 10 µg/ml | 500 µl | 10 ng/ml |
| TGF-β | Peprotech 100-21-100 | 2 µg/ml | 500 µl | 2 ng/ml |

**Supplemental table 4.** FSB medium.

| **FSB medium** |  |  |  |  |
| --- | --- | --- | --- | --- |
| **Component** | **Source/product code** | **Stock concentration** | **Volume/amount** | **Final concentration** |
| CDM/PVA | N/A | N/A | 500 ml | N/A |
| FGF-2 | Qkine Qk027 | 4 µg/ml | 1.5 ml | 12 ng/ml |
| SB 431542 | Apexbio<br>A8249-APE-10mg | 20 mM<br>in DMSO | 250 µl | 10 nM |

**Supplemental table 5.** MEF medium.

| **MEF medium** |  |  |  |  |
| --- | --- | --- | --- | --- |
| **Component** | **Source/product code** | **Stock concentration** | **Volume/amount** | **Final concentration** |
| DMEM/F12 | Thermofisher<br>31330-038 | N/A | 450 ml | N/A |
| FBS | Merck/SigmaAldrich<br>F9665-500ML | N/A | 50 ml | 10% |
| Pen/Strep | Thermofisher<br>15140122 | 100x | 5 ml | 1x |
| β-mercaptoethanol | Merck/SigmaAldrich<br>M3148-25ML | 100 mM | 0.5 ml | 0.1 mM |

**Supplemental table 6.** KSR medium.

| **KSR medium** |  |  |  |  |
| --- | --- | --- | --- | --- |
| **Component** | **Source/product code** | **Stock concentration** | **Volume/amount** | **Final concentration** |
| Advanced DMEM/F12 | Thermofisher<br>12634-028 | N/A | 400 ml | N/A |
| KnockOut Serum Replacement | Thermofisher<br>A3181502 | N/A | 100 ml | 20% |
| L-Glutamine | Thermofisher<br>25030032 | 200 mM | 5 ml | 2 mM |
| Pen/Strep | Thermofisher<br>15140122 | 100x | 5 ml | 1x |
| β-mercaptoethanol | Merck/SigmaAldrich<br>M3148-25ML | 100 mM | 0.5 ml | 0.1 mM |
| **Complete KSR medium (KSRC):** |  |  |  |  |
| FGF-2 | Qkine Qk027 | 4 µg/ml | 3.125 ml | 12 ng/ml |

**Supplemental table 7.** Primary antibodies for ICC.

| Neural crest cells |  |  |
| --- | --- | --- |
| Antibody | Source/product code | Dilution |
| Anti-TFAP2a/AP2-a<br>(rabbit monoclonal) | Abcam Ab108311 | 1/200 |
| Anti-p75/NGFR<br>(mouse monoclonal) | Santa Cruz Biotechnology sc-271708 | 1/200 |
| Smooth muscle cell |  |  |
| Antibody | Source/product code | Dilution |
| Anti-calponin/CNN<br>(mouse monoclonal) | Santa Cruz Biotechnology sc-58707 | 1/250 |
| Anti-calponin/CNN<br>(mouse monoclonal) | Merck/SigmaAldrich C2687 | 1/250 |
| Anti-transgelin/TAGLN/SM22a<br>(rabbit polyclonal) | Abcam ab14106 | 1/500 |

**Supplemental table 8.** Directly conjugated and primary antibodies for flow cytometry.

| Antibody | Source/product code | Dilution |
| --- | --- | --- |
| Neural crest cells |  |  |
| Isotype control, 488nm | Santa Cruz Biotechnology sc-3890(AF488) | 1/10 |
| Isotype control, 647nm | Santa Cruz Biotechnology sc3890(AF647) | 1/10 |
| Antri-TFAP2a/AP2-a | Santa Cruz Biotechnology sc-12726-AF488 | 1/10 |
| Anti-p75/NGFR-AF647 | Santa Cruz Biotechnology sc-271708-AF647 | 1/10 |
| Smooth muscle cells |  |  |
| Isotype control, 488nm | Santa Cruz Biotechnology sc-3890(AF488) | 1/10 |
| Isotype control, 647nm | Santa Cruz Biotechnology sc3890(AF647) | 1/10 |
| Anti-calponin/CNN (mouse monoclonal) | Santa Cruz Biotechnology sc-58707-AF647 | 1/10 |
| Anti-transgelin/TAGLN/SM22a<br>(rabbit polyclonal) | Santa Cruz Biotechnology sc-53932-AF488 | 1/10 |

**Supplemental table 9.**
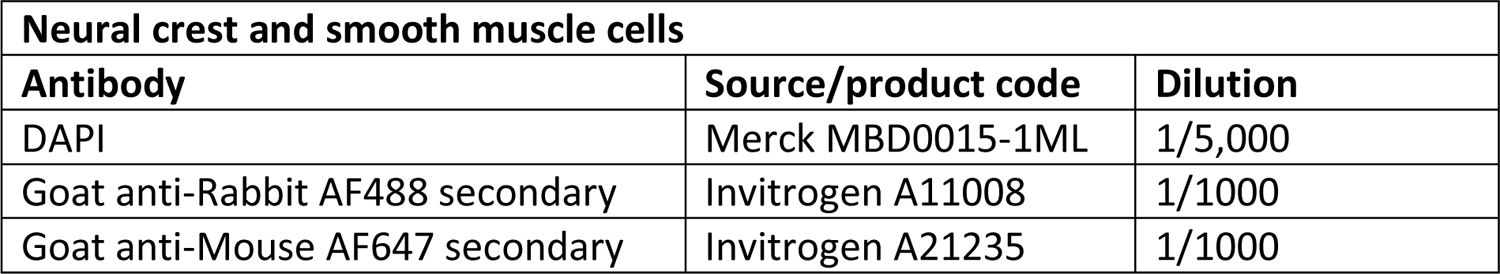
Secondary antibodies and DAPI for ICC.

| Neural crest and smooth muscle cells |  |  |
| --- | --- | --- |
| Antibody | Source/product code | Dilution |
| DAPI | Merck MBD0015-1ML | 1/5,000 |
| Goat anti-Rabbit AF488 secondary | Invitrogen A11008 | 1/1000 |
| Goat anti-Mouse AF647 secondary | Invitrogen A21235 | 1/1000 |

## Reagents

### Collagenase

Collagenase medium: 400 ml Advanced DMEM/F12, 100 ml KnockOut Serum Replacement. 0.5 g Collagenase (Fisher Scientific 17104019) was dissolved in 500 ml collagenase medium, sterile filtered and stored in 50 ml aliquots at −20°C.

**DABCO:** 2.5% DABCO in 90% glycerol with 1.25mM Tris pH 8.0

**PBE Buffer:** PBS with 2 mM ETDTA and 0.5% BS

**Flow cytometry buffer:** PBS with 0.2% Saponin, 5% FBS and 2 mM EDTA

**Neural crest ICC buffers:** PBS with 0.2% Saponin and 5% FBS

PBS with 20% FBS blocking buffer

**SMC ICC buffers:** PBS with 0.1% Triton X-100

PBS with 5% BSA blocking buffer

PBS with 1% BSA otherwise

---

**Supplemental table 10.** Reagents.

| Item | Source/product code | Comment |
| --- | --- | --- |
| Bovine serum albumin (BSA) | Europa Bioproducts EQBAH |  |
| Collagenase | Fisher Scientific 17104019 |  |
| DABCO | Merck/SigmaAldrich D27802-100G |  |
| EDTA | Fisher Scientific BP2482-500 |  |
| Gelatin | Merck/SigmaAldrich G1890-500G |  |
| Glycerol | Fisher Scientific G/0650/17 |  |
| MEF cells, irradiated | Thermofisher A34181 |  |
| PBS (no Ca, no Mg) | Thermofisher 14190-094 |  |
| PFA (4%) in PBS | Alfa Aesar J61899.AP |  |
| ROCK Inhibitor Y-27632 dihydrochloride 10mg | Apex Bio A3008-APE-10mg | 10 mM stock solution in nuclease-free water |
| Saponin | Alfa Aesar A18820.14 |  |
| Stem Cell Banker DMSO free | AMSBIO 11890F |  |
| Tris base | Merck/SigmaAldrich T6066-1KG |  |
| Triton X-100 | Merck/SigmaAldrich T9284-100ML |  |
| TrypLE | Thermofisher 12604-021 |  |
| Vitronectin-XF | Stem Cell Technologies #07180 |  |

**Supplemental table 10.** Cell lines.

| Item | Source/product code | Comment |
| --- | --- | --- |
| MEF cells, irradiated | Thermofisher A34181 |  |
| H9 human stem cells | Wicell |  |
| HPSi0114i-eipl_1 | HipSci HPSi0114_xxxx | n fem |
| HPSi0214i-wibj_2 | HipSci HPSi0214i_wibj_2 | n fem |
| HPSi0314i-sojd_3 | HipSci HPSi0314i_sojd_3 | n fem |
| HPSi0414i-seru_7 | HipSci HPSi0414_xxxx | n fem |

